# QuickRNASeq: Guide for Pipeline Implementation and for Interactive Results Visualization

**DOI:** 10.1101/125856

**Authors:** Wen He, Shanrong Zhao, Chi Zhang, Michael S. Vincent, Baohong Zhang

## Abstract

Sequencing of transcribed RNA molecules (RNA-seq) has been used wildly for studying cell transcriptomes in bulk or at the single-cell level (1, 2, 3) and is becoming the *de facto* technology for investigating gene expression level changes in various biological conditions, on the time course, and under drug treatments. Furthermore, RNA-Seq data helped identify fusion genes that are related to certain cancers (4). Differential gene expression before and after drug treatments provides insights to mechanism of action, pharmacodynamics of the drugs, and safety concerns (5). Because each RNA-seq run generates tens to hundreds of millions of short reads with size ranging from 50bp-200bp, a tool that deciphers these short reads to an integrated and digestible analysis report is in high demand. QuickRNASeq (6) is an application for large-scale RNA-seq data analysis and real-time interactive visualization of complex data sets. This application automates the use of several of the best open-source tools to efficiently generate user friendly, easy to share, and ready to publish report. Figure 1 illustrates some of the interactive plots produced by QuickRNASeq. The visualization features of the application have been further improved since its first publication in early 2016. The original QuickRNASeq publication (6) provided details of background, software selection, and implementation. Here, we outline the steps required to implement QuickRNASeq in user’s own environment, as well as demonstrate some basic yet powerful utilities of the advanced interactive visualization modules in the report.

## 1 Introduction

Since its publication in early 2016, the QuickRNASeq pipeline has been adopted by many bioinformatics scientists and experimental researchers to do RNA-seq data analysis, for its expedient automation of the analysis pipeline and its convenient visualization. This detailed protocol provides instructions on installing every component of the pipeline, preparing sample data, running the pipeline for individual sequencing runs, merging results from different runs, interpreting the outcome and making figures for visualization. The goal of this protocol is to show you how to get the QucikRNASeq report from fastq files, as well as how to use the visualization features of the report. The structure of this protocol is outlined as follows. Section 2 is about Materials, and it describes the required hardware, software, and reference genome. Section 3 is on Methods. Section 3.1 describes the input files. Section 3.2 describes the command line call for individual runs. Section 3.3 is for combining the results from 3.2 to summary files and generating the report. Section 3.4 describes how to explore the report and make various plots from the interactive visualization tools. And finally, section 4 includes notes for more productive use of QuickRNASeq.

## 2 Materials

### 2.1 Hardware

The QuickRNASeq package is fully tested on an HPC cluster using the IBM Platform LSF (Load Sharing Facility) or on a standalone workstation running Linux. Since the mapping step of millions of reads is a memory-demanding procedure. It is recommended to have 64GB per running instance. Other required hardware includes storage arrays with a high I/O throughput such as EMC Isilon if hundreds of samples are processed at the same time in parallel.

### 2.2 Software prerequisites

Many open-source tools developed for RNA-seq data analyses were tested before QuickRNASeq settled on the following 5 applications. STAR (7) was chosen for read alignment, or mapping, to reference genome and transcriptome assembly. FeatureCounts (8) from Subread package was adopted for counting reads to genomic features such as genes, exons, promoters and genomic bins. VarScan (9) was used for variant calling. RSeQC (10) was chosen for RNA-seq quality control. Samtools (11) provides various utilities for manipulating alignments in the SAM format. These open source tools should be installed as instructed below. Names of directories are for demonstration only, which should be replaced by your own names.

#### 2.2.1 STAR

Download STAR from https://github.com/alexdobin/STAR/releases Install STAR to /opt/ngsapp/STAR_2.4.0k/bin/Linux_x86_64

#### 2.2.2 Subread

Download Subread packages from http://subread.sourceforge.net/ Install Subread to /opt/ngsapp/subread-1.4.6/bin

#### 2.2.3 VarScan

Download JAR file from http://varscan.sourceforge.net/ Install VarScan to /opt/ngsapp/bin/VarScan.v2.4.0.jar

#### 2.2.4 RSeQC

Download and install RSeQC from http://rseqc.sourceforge.net/ Install RSeQC to /opt/ngsapp/anaconda/bin

#### 2.2.5 Samtools

Download Samtools from http://sourceforge.net/projects/samtools/files/ Install Samtools to /opt/ngsapp/bin.

### 2.3 Download QuickRNASeq package

QuickRNASeq (5) is available from sourceforge. Follow this link to download the source code: https://sourceforge.net/projects/quickrnaseq. The protocol is based on version 1.2.

We have QuickRNASeq installed at directory /opt/ngsapp/QuickRNASeq.

### 2.4 Preparation of genome fasta file, annotation and index

Here we show how to create the index for RNA-seq analysis using Gencode release 23 of human genome GRCh38 as an example. For details of STAR related command line parameters, please refer to recent publication from Dobin and Gingeras on optimizing RNA-seq mapping with STAR (12).

1. Download genome fasta file ftp://ftp.sanger.ac.uk/pub/gencode/Gencode_human/release_23/GRCh38.primary_assembly.geno_me.fa.gz
2. Download gene annotation in GTF format ftp://ftp.sanger.ac.uk/pub/gencode/Gencode_human/release_23/gencode.v23.annotation.gtf.gz
3. Unzip and rename genome and GTF file Unzip and rename genome fasta file and GTF file as GRCh38.primary.genome.fa, GRCh38.gencode.v23.gtf respectively. Save these two files in the corresponding project data folder. In this example, genome fasta file is saved to directory /opt/fasta; GTF annotation file to directory /opt/gencode
4. Prepare annotation and BED files using utility functions in QuickRNASeq
5. Make sure you are in directory /opt/gencode, and call QuickRNASeq utility functions as shown below:

~~~
*/opt/ngsapp/QuickRNASeq/gtf2bed.pl GRCh38.gencode.v23.gtf > GRCh38.gencode.v23.bed
/opt/ngsapp/QuickRNASeq/gtf2annot.pl GRCh38.gencode.v23.gtf > GRCh38.gencode.v23.annot
/opt/ngsapp/bin/samtools faidx GRCh38.primary.genome.fa*
~~~
6. Create genome index file
7. In this example, we are creating a genome index for read length up to 100bp. Under directory /opt/STAR/, create a directory called GRCh38_gencode23_100. Move to this GRCh38_gencode23_100 directory, and create genome index as shown below. The option sjdbOverhang is set at 99. In general, sjdbOverhang is set as “read length - 1”. An example command is listed below:

~~~
*/opt/ngsapp/STAR_2.4.0k/bin/Linux_x86_64/STAR ‐‐runThreadN 32 ‐‐runMode genomeGenerate ‐‐
genomeDir /opt/STAR/GRCh38_gencode23_100 –genomeFastaFiles
/opt/fasta/GRCh38.primary.genome.fa–sjdbGTFfile /opt/gencode/GRCh38.gencode.v23.gtf ‐‐
sjdbOverhang 99*
~~~
8. Find the chromosome where MHC genes are located
9. MHC genes are highly polymorphic, which makes this region ideal for checking sample SNP concordance. In the human genome, the MHC region occurs on chromosome 6. Get the corresponding coordinate for chromosome 6 from chrNameLength.txt file in the STAR index result folder. In this case, the coordinate is 1-170805979.
10. After the above steps are completed, you are ready to set the reference genome related parameters in the configuration file. Refer to section 3.1.3 part 6 for instructions.

## 3 Methods

The QuickRNASeq pipeline can be divided into the following 3 main steps:

1. Prepare RNA-seq input data, including a sample configuration file
2. Process individual samples, including mapping, counting and QC
3. Merge results from individual sample and generate an integrated report

The first step is specific to individual runs and samples within those runs. This step needs to be tailored for each RNA-seq run. The last step is more or less fixed. A master-cmd.sh file included in the package contains the common commands to be called for step 2 and step 3. Nevertheless, all environmental variables need to be set correctly to ensure the scripts in master-cmd.sh will work well. All these steps should be performed under a project folder for all samples belonging to a specific project.

After downloading QucikRNASeq1.2, you will see a directory named “test_run”. This is an example project directory. We are using the same 48 GTEx samples from 5 donors as in the original QuickRNASeq publication (6). This test_run project directory contains key files for running QuickRNASeq. The following discussions describe the contents of these files, and step-by-step instructions to guide you through the process.

### 3.1 Prepare RNA-seq input data

#### 3.1.1 Prepare a sample annotation file and a sample ID file

1. Annotation file

To run QuickRNASeq, a user needs to provide meaningful annotations for all samples. A proper annotation file should be in tab delimited text format. The first and second columns correspond to sample and subject identifiers, respectively. Although not required, it is highly recommended to use “sample_id” and “subject_id” for the first two columns while the rest of the columns are flexible, based on project design. The sample.annotation.txt file in test_run directory has columns as “Run”, “subject_id”, “histological_type”, and “sex”.

2. Sample ID file

Sample ID file contains one unique sample ID per line. There is no column header. The allIDs.txt file in test_run directory lists all 48 samples in this demo project. For example, the first sample ID is “SRR607214”.

#### 3.1.2 Prepare fastq files for each individual sample

For paired end sequencing, prepare two fastq files, one for each read. Format will be sample_id_1.fastq.gz and sample_id_2.fastq.gz. For example, sample SRR607214 will have two files: SRR607214_1.fastq.gz, and SRR607214_2.fastq.gz.

Some new Illumina sequencing platforms, such as NextSeq500, generate 8 files as output for each sample in paired end sequencing. In this case, we need to concatenate these fastq files into 2 files, one for each read of paired end sequencing. Make sure the concatenation order is the same for both files.

There will be only one fastq file per sample if the sequence run is single end.

At the end of this step, we will have “N” numbers of fastq files if the run contains “N” single read samples. Or we have “2xN” of fastq files if the sequencing is paired end run. We save these files in a directory called fastq.

#### 3.1.3 Set up run configuration file

File run.config is a project-specific configuration file that contains all sequencing, genome, and software related information for QuickRNASeq analysis. Genome and software portions only need to be changed if there are updates on tools or alterations on genome, index and/or annotation. The sequencing run-specific portion is what we need to modify for each analysis.

Please refer to $QuickRNASeq/star-fc-qc.config.template for more details. You can copy star-fc-qc.config.template in QuickRNASeq package to your project folder and then customize it to your environment. Please see directory test_run for an example of run.config file.

##### 1. Set FASTQ_DIR

FASTQ_DIR is the directory where the fastq files are located. You can set a fastq directory within the project folder to store all fastq files, or you can store your fastq file in another location.

##### 2. Set the suffix for fastq file

QuickRNASeq will automatically add “_1. FASTQ_SUFFIX” and “_2. FASTQ_SUFFIX” to each sample ID in the allID.txt file, and look for these files in the FASTQ_DIR. The name of the fastq file should match the name in the allID.txt file. For example, for sample SRR607214, if you set FASTQ_SUFFIX=fastq.gz, the program will go to directory FASTQ_DIR and look files SRR607214_1.fastq.gz and SRR607214_2.fastq.gz. Sometimes, the fastq files end as fq.gz, sometimes, it ends as fastq.gz. In our example run.config file, it was set as FASTQ_SUFFIX=fastq.gz.

##### 3. Set strand information

~~~
*STRAND=0 for non-stranded RNA-seq
STRAND=1 for first read forward strand
STRAND=2 for first read reverse strand, for instance Illumina ’s sequencing kit*
~~~

##### 4. Set sequencing depth

There are two choices for sequencing depth option. Set it to “regular” if the sequencing run generates 40-80 million reads; or set it to “deep” if the run generates 100 million reads or more. For example: SEQUENCE_DEPTH=regular.

##### 5. Set sequence type

This is to state whether your read is paired or single. e.g. SEQUENCE_TYPE=pair. 2.4

##### 6. Set species-specific genome index and GTF file

These options will not change unless the genome reference changes. Please refer to section 2.4 for instruction on how to generate these species-specific files.

~~~
*GENOME_FASTA=/opt/fasta/GRCh38.primary.genome.fa
GENOME_INDEX=/opt/STAR/GRCh38_gencode23 100
GENOME_ANNOTATION=/opt/gencode/GRCh38.gencode.v23.annot
GTF_FILE=/opt/gencode/GRCh38.gencode.v23.gtf
BEDFILE=/opt/gencode/GRCh38.gencode.v23.bed
CHR_REGION=chr6:1-170805979*
~~~

##### 7. Set the environmental variables for tools installed at step 2.2

Software locations will remain the same unless there is a major update.

~~~
*STAR_RNA =/opt/ngsapp/STAR_2.4.0k/bin/Linux_x86_64
FEATURECOUNTS=/opt/ngsapp/subread-1.4.6/bin
RSeQC=/opt/ngsapp/anaconda/bin
VARSCAN_JAR=/opt/ngsapp/bin/VarScan.v2.4.0.jar
SAMTOOLS=/opt/ngsapp/bin*
~~~

### 3.2 Run the QuickRNASeq script to process individual samples

QuickRNASeq calls R and Rscript. Please make sure R version 3.1 or above and these R packages, ggplot2, edgeR, scales, and reshape2 are installed on your machine.

Under your project folder, invoke mapping, counting, QC, and SNP call for each sample by calling star-fc-qc.sh. Make sure that the $PATH environmental variable includes the path to QuickRNASeq_1.2 location. Because this step is computationally intensive, it is advised to run this command on HPC clusters using LSF as a job scheduler. A separate result folder will be created for each sample under the project folder. In addition to LSF, there is a list of notable job scheduling software available to choose from. For a cluster using a job scheduler other than LSF, star-fc-qc.sh needs to be twisted or modified. For people who have no access to a HPC cluster, we offer star-fc-qc.ws.sh, a customized script working in a standard Linux workstation. Of course, analyzing a large RNA-seq dataset in a single workstation is not typical. Below is the command call example.

~~~
     *# ENVIRONMENT
     export QuickRNASeq={QuickRNASeq_installation_Directory}
     # e.g. export QuickRNASeq=/opt/ngsapp/QuickRNASeq_1.2
     export PATH=$QuickRNASeq:$PATH
     star-fc-qc.sh allIDs.txt run.config

     #run the following command if you run the analysis on a standalone workstation
     #star-fc-qc.ws.sh allIDs.txt run.config*
~~~

### 3.3 Merge results from individual samples and generate an integrated report

As in the previous steps, this step also runs under the project directory. We run the merging and summarization step when all jobs are finished for each sample. The sample.annotation.txt should include all samples to be merged. Each sample has to be processed as listed in step 3.2. Below are commands to run in order to generate the report. “GENE_ANNOTATION” points to a file containing gene descriptions that can be obtained by running “Rscript $QuickRNASeq /QuickRNASeq_html/getEnsemblAnno.R”.

~~~
*#Summarization, only run it when all jobs are finished in the first step
     export GENOME_ANNOTATION=/opt/gencode/hg19.gencode.v19.annot
     export GENE_ANNOTATION=/opt/gencode/Ensembl_v75_hg19_Gencode.v19_human.txt.gz
     nohup star-fc-qc.summary.sh sample.annotation.txt &> Results.log
     #run the command line below if you run the analysis in a workstation
     #star-fc.summary.sh sample.annotation.txt*
~~~

#### 3.3.1 QuickRNASeq test run

We made a test run available for you to test the QuickRNASeq software before applying this tool to your data. Adjust QuickRNASeq to point to your QuickRNASeq installation folder. Please refer to $QuickRNASeq/test_run folder for:

- allIDs.txt: sample identifiers
- sample.annotation.txt: annotation file
- run.config: sample configuration file
- master-cmd.sh: command lines for test runs. Please run step #2 after step #1 finishes.

#### 3.3.2 Description of output files

The output of the merge command is a directory called Results. Seven html files (gex.html, index.html*, longitudinal.html, qc_fc-counting-summary.html, qc_expr_count_RPKM.html, qc_overview.html, qc_star-mapping-summary.html) and three directories (package, QC, summary) will be generated within the Results directory.

This Results directory can be copied to your laptop or desktop. Open index.html within Results directory to access the interactive report in html format. Alternatively, this Results directory can be hosted on a web sever to share with other group members, which also makes this QuickRNASeq report available at all times. Summary directory contains all summary files which are displayed on the html report under the “Raw Data Files” section.

### 3.4 Explore integrated and interactive report

Open the index.html file under Results directory and you will have access to all data and figures. You will be able to drill down RNA-seq analysis results in an interactive way. We implemented the interactive data visualization in QuickRNASeq using these JavaScript-based open-source libraries including JQuery (12), D3 (Data-Driven Documents) (13), canvasXpress (14), SlickGrid (15), and Nozzle (16). The figures and tables in QC Metrics and QC Plot portion of the QuickRNASeq web report have been introduced and described in the original publication (6). Some of these figures are showcased in Figure 1. Below, we focus on the interactive plotting features that can be accessed by clicking the pointing hand next to “RPKM Values on Genes” under the Expression Table section.

**Fig. 1.**
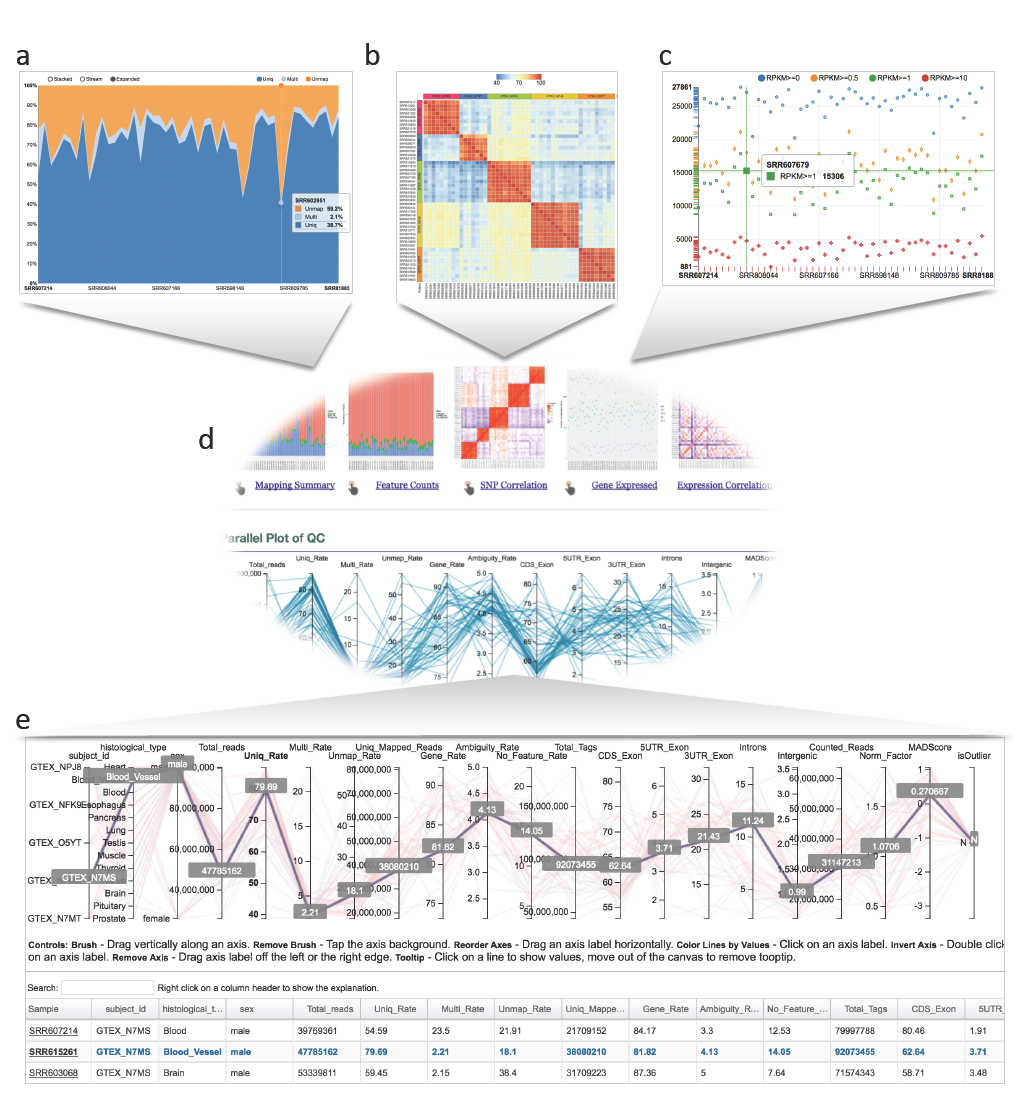
Interactive plots from QuickRNASeq report. Figures (a, b, c) can be retrieved by clicking on the pointing hands as shown in figure 1d. On any of these interactive plots, mouse over each sample displays associated sample QC metrics, (a) Read mapping summary in the expanded display mode, (b) SNP concordance matrix of 48 samples from 5 donors. Samples from the same donor should be highly concordant, (c) Gene expression chart, which shows the number of genes past various expression thresholds, (d) Center portion of the QuickRNASeq report, (e) Parallel plot linking multiple QC measures for the same samples plus table of multi-dimensional QC measures.

#### 3.4.1 Meaning of mouse icons

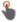Pointing hand, click to get interactive plot 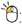 Left click;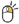Right click;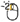Double left click; 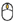Scroll middle wheel

#### 3.4.2 Create a boxplot for one gene

1. Figure 2 combines 6 charts to demonstrate how to create a boxplot for gene expression of a single gene. From the main QuickRNASeq report HTML page, click on the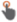 “pointing hand” icon in the Expression Table section to get to gene expression table. A new HTML page will show up, see Fig. 2a. This webpage is from file gex.html, which was generated as a result of running star-fc-qc.summary.sh.
2. Search by keyword and then left click on any column except the first two on a gene (Fig. 2a). As demonstrated in Fig. 2a, we searched by “kinase”, then selected gene “CAMKK1”.
3. A new window pops up which displays a dot plot of gene expression level in RPKM value for kinase CAMKK1 (Fig. 2b). Please note that the X-axis is smaple ID.
4. Right click on any plot area to bring up the drop down menu for sample grouping, data transformation and chart customization. As shown in Fig 2c, samples were grouped by following menu “Group Samples” and then “histological_type” for box plot. Please note that in Fig. 2c, X-axis is histological_type. Click on any plot area to hide the menu and you should see the boxplot (not shown here). The sample features are gathered from the “sample.annotation.txt” file.
5. Data can be transformed to various scales by right clicking on the plot to bring up the menu and then following “Data” -> “Transform” -> “Log Base 2” for log2 transformation (Fig. 2d).
6. User can also adjust the font of the sample label, add Y-axis, change window and canvas size, color data points, and explore many other visualization features, see Note 4.1.
7. Data points can be connected as shown in Fig. 2e by subject_id.
8. When you are satisfied with the settings, move mouse up to the top of the canvas to activate the top menu, where you will see a “Camera” icon. Click on the “Camera” icon to get a screenshot in png format for publication (Fig. 2f).

**Fig. 2.**
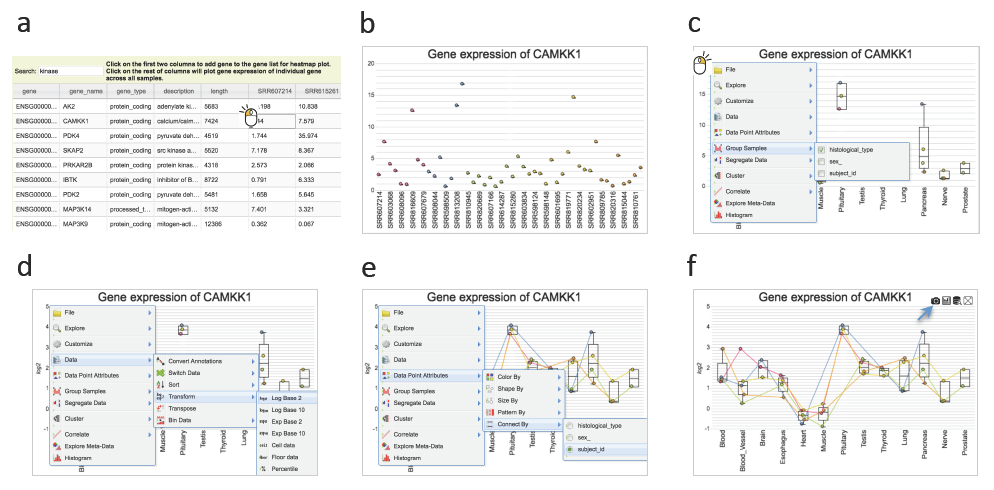
Boxplot of a gene, (a) From gene expression table, search genes by keyword “kinase”, and select gene CAMKK1. (b) Dot plot of CAMKK1 gene expression across all samples, (c) Group sample by histological type, (d) Log2 transformation of expression level, (e) Connect data point by subject identifier, (f) Take the screenshot of the boxplot as a png image.

#### 3.4.3 Create a heatmap for multiple genes

1. Figure 3 consists of several charts which illustrate the steps involved for drawing a gene expression heat map. As shown in Fig. 3a, in the large text box below the table of expression, there are 10 genes, which are “TNFRSF1B CDA FAM131C BAI2 C1orf170 LRRC38 FCN3 C4BPA NPPB PLA2G5”. These genes can be typed in the text box, or select one by one from the expression table above the text box. You can also copy and paste the gene list separated by space or comma. Copying and pasting the gene list from an Excel file also works. It is suggested to enter the official HUGO gene symbol for each gene. After you have the gene list, click on “Plot Heatmap” button, a heatmap shows up, as in Fig. 3b.
2. Expression level from different genes could vary widely. It is a good idea to have the heatmap displays gene expression level in log2 format. Right click on any plot area to bring up the drop down menu for data transformation. The example in Fig. 3c is transforming data into Log Base 2.
3. Cluster samples by following menu “Cluster” -> “Cluster Samples”. Cluster variables by following menu “Cluster” -> “Cluster Variables”. See Fig. 3d for a demonstration.
4. Move mouse up to the top of the canvas to activate the top menu, where you will see the “Camera” icon again, and then click on the “Camera” icon to get a png image ready for publication (Fig. 3e).
5. Figure 3f is for expression correlation among the genes in the gene list. Click “Plot Correlation” to get the correlation picture shown in Fig. 3f. Double clicking on any square will show the correlation value between two genes. The example in Fig. 3f shows the expression correlation across samples between gene LRRC38 and gene C1ORF170 is 0.863.

**Fig. 3.**
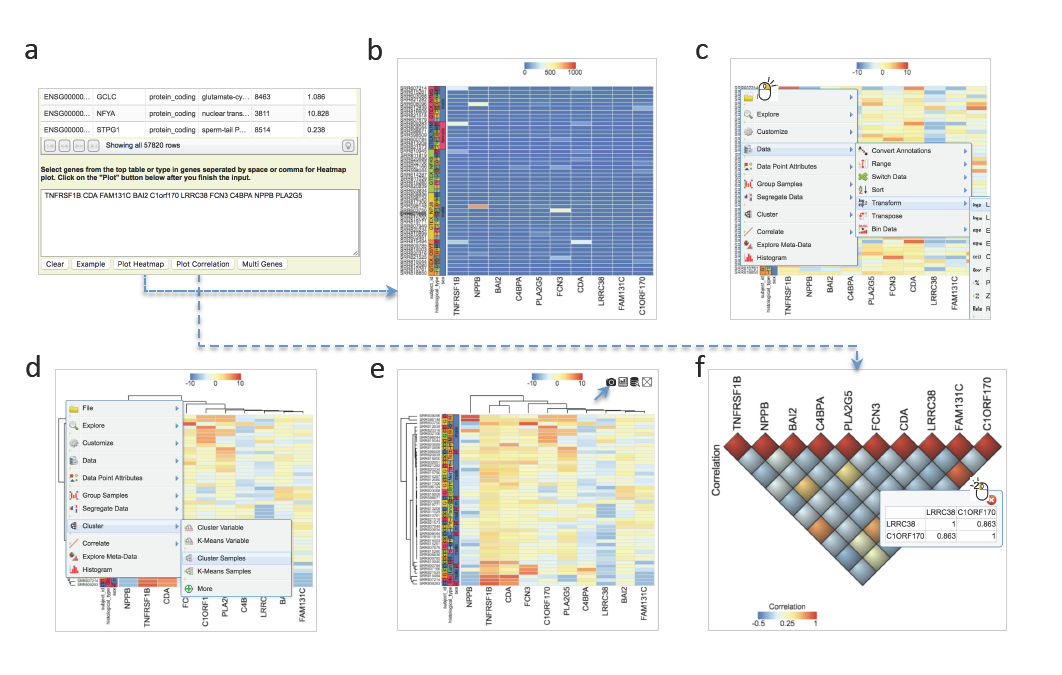
Generation of heatmap for a list of genes, **(a)** Select from the above table or enter a list of genes into the text box. **(b)** Initial heatmap. **(c)** Log2 transformation of expression level, **(d)** Cluster samples and variables, (e) A final heatmap ready to be saved in png format, **(f)** Gene expression correlation plot.

### 3.5 Make the report publicly available at github.com

#### 3.5.1 Create a repository by login github.com

You will need a GitHub account to perform this step. If you don’t have a GitHub account, create one first by going to https://github.com. To create a GitHub repository, please follow these two steps.

1. Click on the “New repository” icon as shown in Figure 4a after GitHub account login.
2. Type in project name and description and then click “Create repository” icon as illustrated in Figure 4b. Please use your own project name instead of “RNASeq_1” that is for illustration purpose only.

**Fig. 4.**
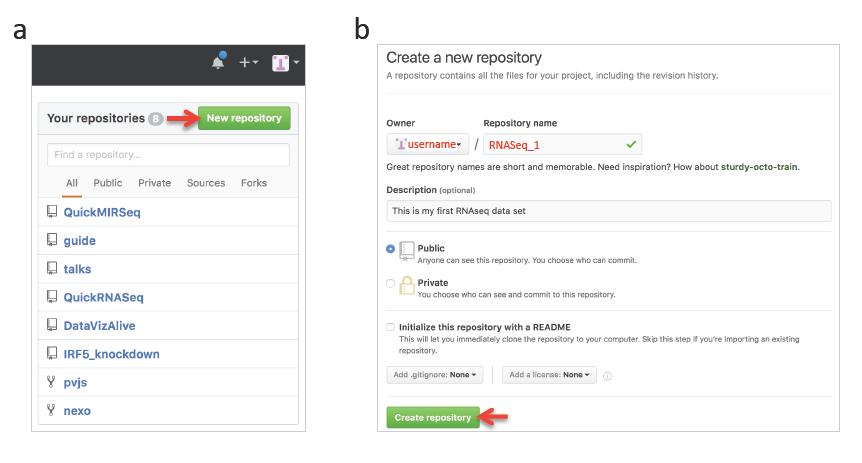
Creation of a GitHub repository, (a) Repository panel is located on the right side of the web page after GitHub login. The red arrow points to the “New repository” button, (b) Fill in “Repository name” field using your own name and check the “Public” radio button if you want to publish it to the Internet then click “Create repository” to finish.

#### 3.5.2 Commands to publish the report files to GitHub repository

Change texts in red to your own settings.

~~~
     *git clone https://github.com/username/RNASeq_1.git
     cd RNASeq_1
     git checkout ‐‐orphan gh-pages
     cp -R path2result/Results/*.
     git add .
     git commit -a -m “Adding RNASeq_1 results from QuickRNASeq”
     git push origin gh-pages*
~~~

Now, the report should be available at http://username.github.io/RNASeq_1

The demo page for the example data set is at http://baohongz.github.io/QuickRNASeq

## 4 Notes

### 4.1 Further customization of plots

1. The font of sample labels will be enlarged by following “Customize” -> “Sample Labels” -> “Font” -> “Bigger”. The more you click on the “Bigger” button, the larger the font becomes.
2. Add y-axis title by following “Customize” -> “Axes Titles” -> “Text”. Type in title in the input box and then click the nearby cycling button.
3. The size of Pop-up Window and Canvas can be altered by click-and-drag the left bottom corner as indicated by the black arrow that will appear while the mouse moves over.
4. Follow “Data Points Attributes” -> “Color by” to color data points based on certain feature.

### 4.2 Wired characters in input files

Although we have taken multiple measures to either remove or replace R unfriendly characters and unnecessary blank spaces in user input files such as sample.annotation.txt, it is recommended to use only alphanumeric and tab characters in these files. If Microsoft Excel is used to create the sample.annotation.txt, please make sure you save it in tab-delimited format.

